# Feasibility of real-time *in vivo* ^89^Zr-DFO-labeled CAR T-cell trafficking using PET imaging

**DOI:** 10.1101/789727

**Authors:** Suk Hyun Lee, Hyunsu Soh, Jin Hwa Chung, Eun Hyae Cho, Sang Joo Lee, Ji-Min Ju, Joong Hyuk Shin, Hyori Kim, Seung Jun Oh, Sang-Jin Lee, Junho Chung, Seog-Young Kim, Jin-Sook Ryu

**Affiliations:** Department of Nuclear Medicine, Asan Medical Center, University of Ulsan College of Medicine, Seoul, Republic of Korea; Department of Radiology, Division of Nuclear Medicine, Hallym University Kangnam Sacred Heart Hospital, Hallym University College of Medicine, Seoul, Republic of Korea; Asan Institute for Life Sciences, Asan Medical Center, Seoul, Republic of Korea; Convergence Medicine Research Center, Asan Medical Center, Seoul, Republic of Korea; Research Institute, National Cancer Center, Gyeonggi-do, Republic of Korea; Department of Biomedical Sciences, Seoul National University, Seoul, Republic of Korea

## Abstract

**Introduction:** Chimeric antigen receptor (CAR) T-cells have been developed recently, producing impressive outcomes in patients with hematologic malignancies. However, there is no standardized method for cell trafficking and *in vivo* CAR T-cell monitoring. We assessed the feasibility of real-time *in vivo* ^89^Zr-p-Isothiocyanatobenzyl-desferrioxamine (Df-Bz-NCS, DFO) labeled CAR T-cell trafficking using positron emission tomography (PET).

**Results:** The ^89^Zr-DFO radiolabeling efficiency of Jurkat/CAR and human peripheral blood mononuclear cells (hPBMC)/CAR T-cells was 70–79%, and cell radiolabeling activity was 98.1–103.6 kBq/10^6^ cells. Cell viability after radiolabeling was >95%. Compared with unlabeled cells, cell proliferation was not significantly different during the early period after injection; however, the proliferative capacity decreased over time (*p* = 0.02, day 7 after labeling). IL-2 or IFN-γ secretion was not significantly different between unlabeled and labeled CAR T-cells. PET/magnetic resonance images in the xenograft model showed that most of the ^89^Zr-DFO-labeled Jurkat/CAR T-cells were distributed in the lung (24.4% ± 3.4%ID) and liver (22.9% ± 5.6%ID) by 1 hour after injection. The cells gradually migrated from lung to the liver and spleen by day 1, and remained stably until day 7 (on day 7: lung 3.9% ± 0.3%ID, liver 36.4% ± 2.7%ID, spleen 1.4% ± 0.3%ID). No significant accumulation of labeled cells was identified in tumors. A similar pattern was observed in *ex vivo* biodistributions on day 7 (lung 3.0% ± 1.0%ID, liver 19.8% ± 2.2%ID, spleen 2.3% ± 1.7%ID). ^89^Zr-DFO-labeled hPBMC/CAR T-cells showed the similar distribution on serial PET images as Jurkat/CAR T-cells. The distribution of CAR T-cells was cross-confirmed by flow cytometry, Alu polymerase chain reaction, and immunohistochemistry.

**Conclusion:** Using PET imaging of ^89^Zr-DFO-labeled CAR T-cells, real time *in vivo* cell trafficking is feasible. It can be used to investigate cellular kinetics, initial *in vivo* biodistribution, and the safety profile in future CAR T-cell development.

## Introduction

Given shifting cancer treatment paradigms, chimeric antigen receptor (CAR) T-cell immunotherapy has been developed very rapidly [1,2]. CAR T-cells, which have been studied as immune regulatory cell therapies, harbor fusion proteins with the extracellular scFv domain of an antibody. These proteins recognize the characteristic antigen on the tumor cell surface and the intracellular co-stimulatory domain for T-cell activation. When CAR T-cells grab the antigen on the surface of the tumor cell, a sequential co-stimulatory signal activates the T-cell and triggers the signaling pathway within the cell, thereby allowing the CAR T-cells to kill the tumor cell [3,4]. Moreover, because of its tumor cell-killing activity, CAR T-cells are a “living drug” that can proliferate in the body and kill tumor cells. They have significantly longer action than conventional chemotherapeutics and antibody drugs [5]. CAR T-cell therapy has shown dramatic anti-cancer effects, particularly in clinical trials for patients with hematological malignancies such as refractory B-cell malignancies, after standard treatment [6–8].

Despite its successful use in patients with B-cell malignancies, there is a lack of substantive understanding of CAR T-cells in the human body: 1) a limited effect of CAR T-cells on solid tumors, 2) the trafficking and biodistribution of CAR T-cells, and 3) the targeting efficacy of CAR T-cells that are injected within a patient’s body. To date, there are no available standardized methods for monitoring *in vivo* behaviors and targeting efficacy of injected CAR T-cells. The most common (but limited) techniques used to identify CAR T-cells in the body are flow cytometry, biopsy/immunohistochemistry (IHC), enzyme-linked immunosorbent (ELISpot) or polymerase chain reaction (PCR) [9–12]. Unfortunately, none of these can monitor CAR T-cells within a live body. To optimize the efficacy of CAR T-cell immunotherapy and to predict potential toxicities, it is necessary to develop a noninvasive imaging system that can enable the monitoring of CAR T-cell trafficking in a real-time manner. Image-based data provides a great deal of information concerning actual tracking, targeting patterns, real-time distributions, and *in vivo* maintenance for CAR T-cell therapies.

Additionally, the FDA Guidance for Industry: Preclinical Assessment of Investigational Cellular and Gene Therapy Products (updated 11/2013) acknowledged that the fate of investigational cell therapy, after *in vivo* administration is important for characterizing the product’s activity and safety information. To determine distribution after cell administration, imaging methods such as radioisotope-labeled cells, genetically engineered cells (e.g., green fluorescent protein expression), and nanoparticle-labeled cells (e.g., iron-dextran nanoparticles) are recommended. Unlike conventional drugs, cell therapies must acquire data through *in vitro* experiments to determine pharmacological activities or unrecognized toxicity. Therefore, animal models are generally recommended for evaluating cell therapies because basic information on the initial behavior, organ distribution, and targeting in the body after cell therapy are important. Nuclear medical imaging is a proper method that enables real-time monitoring of cells in the body.

Positron emission tomography (PET) is a diagnostic imaging method that can evaluate metabolic activities in the body by injecting a radioactive tracer as a nuclear medicine functional imaging technique. PET is a unique and important tool for tracking cells in preclinical and clinical studies [13,14]. It can be used for translational research, moving from preclinical to clinical studies because the technology features high sensitivity and spatial resolution. There are two ways to image cells: direct and indirect labeling. This study was designed to monitor CAR T-cells via direct labeling. Direct labeling of cells immediately marks the cells with a radioisotope through covalent bond conjugation. Cell migration and distribution immediately after cell injection can be monitored. Herein we establish a method of direct labeling for CAR T-cells. Especially since CAR T-cells can be manipulated *ex vivo*, it is possible to track the behavior and the distribution of small numbers of radiolabeled cells after *in vitro* labeling.

^89^Zr has a long physical half-life (78.4 hours) and is therefore suitable for tracking the behavior of CAR T-cells in the body. In previous reports, cells were directly labeled using isotopes conjugated with ^89^Zr-oxine or DFO moiety for cell imaging [15–18]. Recently, Weist et al. proposed that ^89^Zr-oxine would be a clinically translatable method for real-time evaluation of cell therapies, especially CAR T-cells [19]. However, Bansal et al. [16] reported that the ^89^Zr-DFO labeling strategy was superior to ^89^Zr-oxine, with increased cell stability and viability. Based on existing preclinical applications, this study aimed to assess the feasibility of real-time trafficking of ^89^Zr-DFO-labeled CAR T-cells using PET imaging.

## Materials and methods

### Study design

After preparation of CAR-expressing Jurkat (Jurkat/CAR) T-cells and CAR T-cells from human peripheral blood mononuclear cells (hPBMC), all cells were radiolabeled with ^89^Zr-DFO. The viability, proliferation ability, and function of ^89^Zr-DFO-labeled cells were evaluated. We completed then PET/magnetic resonance imaging (MRI) after injection of ^89^Zr-DFO-labeled Jurkat/CAR T-cells or CAR T-cells into mice with a xenograft for cell trafficking. The imaging data were compared with *ex vivo* experiments performed with unlabeled Jurkat/CAR T-cells. The animal study scheme is shown in Fig 1.

**Fig 1.**
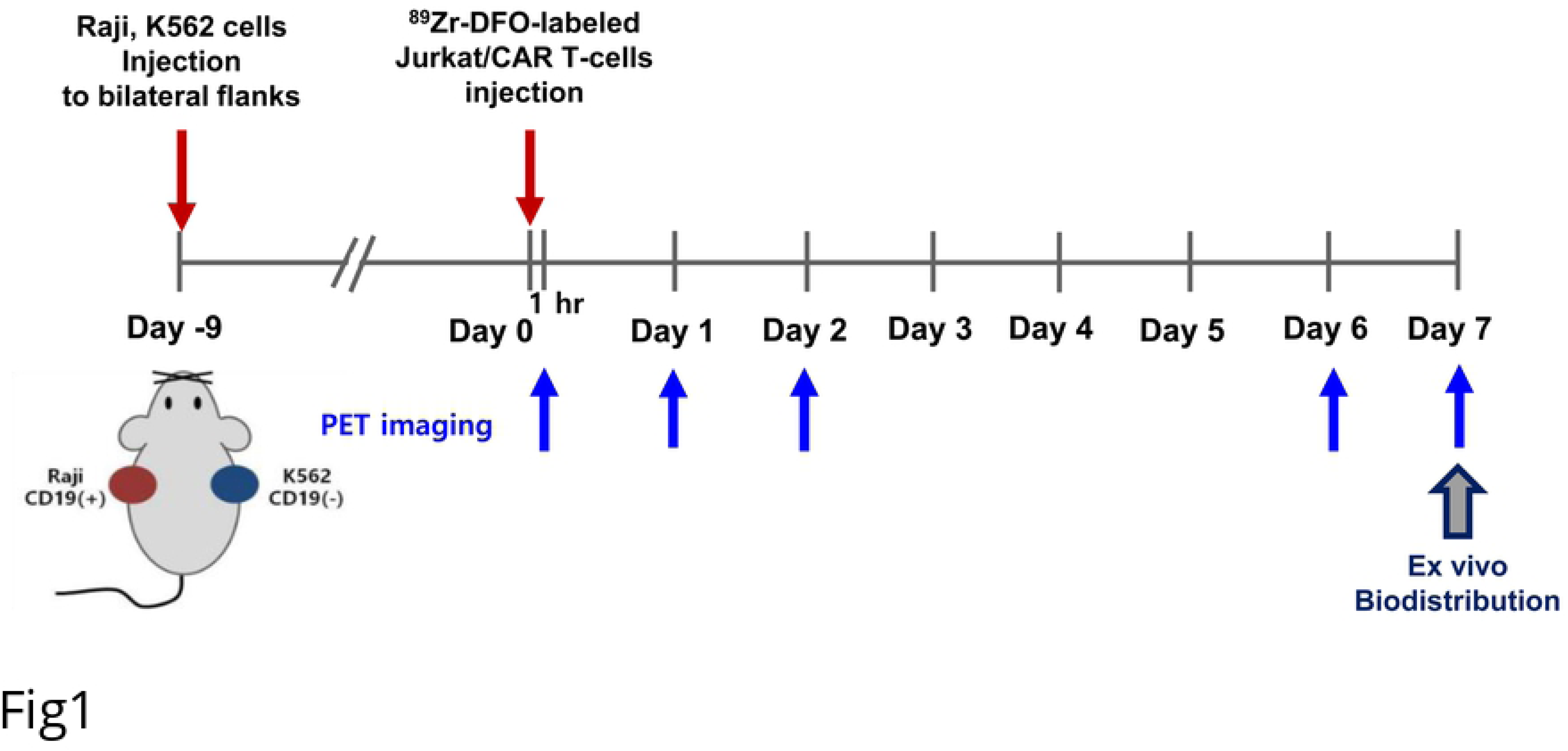
Animal study scheme

The research protocol of this preclinical experimental study with animals was approved by the Institutional Animal Care and Use Committee of the Asan Institute for Life Science (registration no. 2017-12-085). Mice were maintained in accordance with the Institutional Animal Care and Use Committee guidelines of the Asan Institute for Life Science.

### Construction of a lentiviral vector containing CD19-specific CAR

To construct a lentiviral vector encoding CAR specific to CD19, we generated EF1α promoter-driven lentiviral expression vector, pLECE3. pLECE3 was constructed by replacing U6 promoter of pLentiLox3.7 with EF1α promoter along with a few additional cloning sites. The 19BBz consists of anti-CD19 scFv, CD8 Hinge, CD8 transmembrane, 4-1BB, and a CD3 ζ domain, which are identical with Novartis Kymriah product. DNA products of 19BBz domain were amplified by PCR (19BBz; 5’-GATCCgccaccATGGCCTTA CCAGTGA-3’ and 5’-GTTAACttaGCGAGGGGGCAGGGCCTGCAT-3’). The PCR product was then sub-cloned into a pGEM-T-easy vector (Promega, USA). 19BBz was inserted into pLECE3 at the *BamHI/HpaI* site, under the EF1a promoter (Fig 2a).

**Fig 2.**
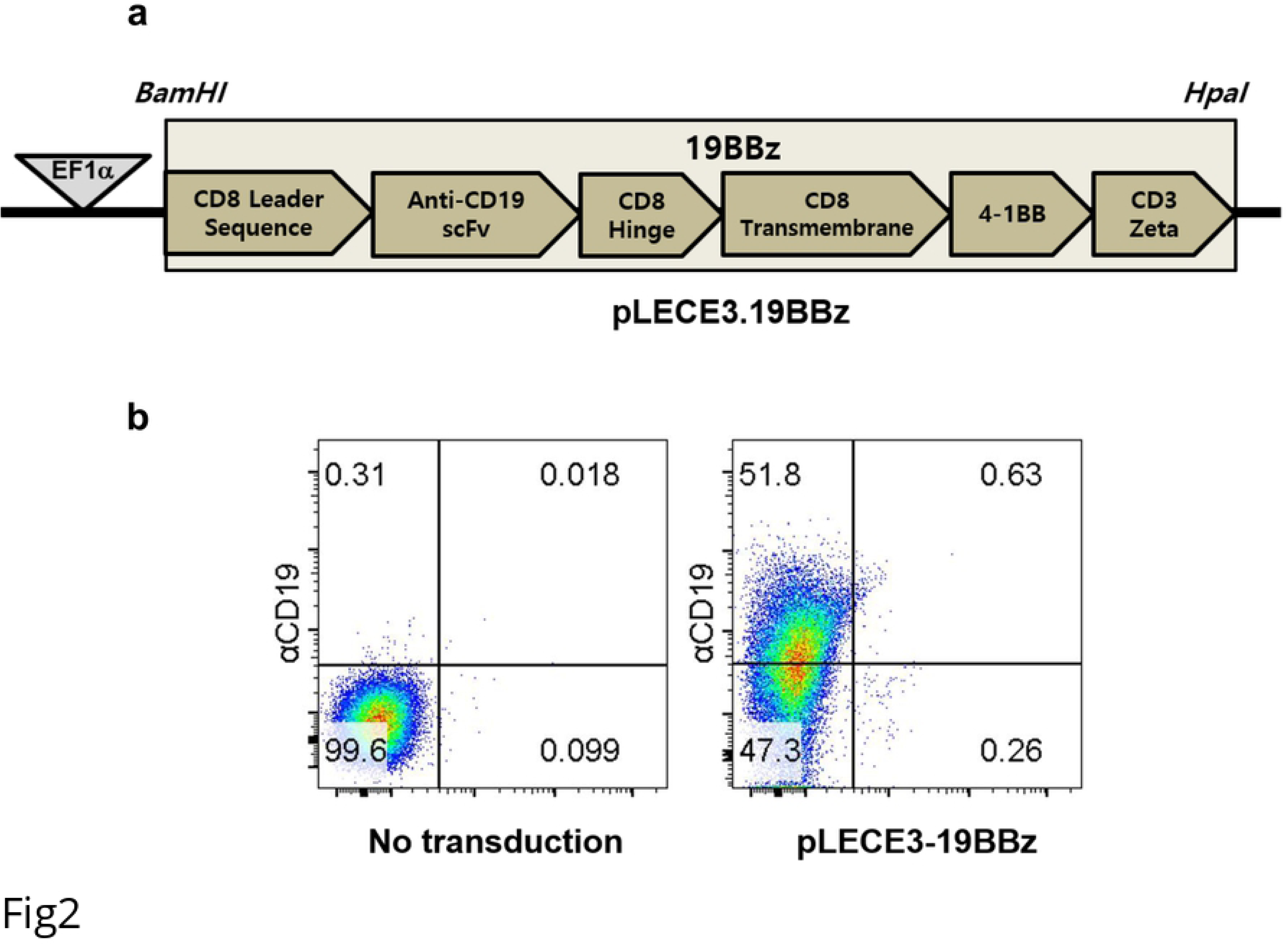
Construction of Jurkat/CAR T-cells specific to CD19. **a** DNA map of pLECE3-19BBz lentivirus vector. **b** CAR expression on Jurkat T-cells. Jurkat T-cells were either left alone (left) or transduced with pLECE-19BBz lentiviruses (right).

### Transfection and lentivirus packaging

The lentiviral vectors were transfected into 293FT cells with Lipofectamine 3000® (Thermo Fisher Scientific), following manufacturer’s protocol. Briefly, 7 × 10^6^ 293T cells were plated in 150 mm petri dishes. The cells were treated with a lipofectamine reagent, viral vectors, and packaging DNA. For high-titer lentiviral purification, cellular supernatants were collected and filtered through a 45 mm pore filter unit (Sartorius AG, Germany) at 48 hours post-transfection, and purified using ultracentrifugation.

### Transduction of lentivirus into Jurkat T-cells

We plated 4 × 10^5^ Jurkat T-cells (ATCC, TIB-152) one day before transduction. The next day, the cells were infected with the lentiviruses (MOI 1000), with polybrene (8 μl/ml, Merk, Germany) added to increase the efficiency of transduction. Then, a 90-minute spin infection was performed (800×g). We sorted transduced Jurkat T-cells expressing 19BBz 72-hours later using FACS Aria (BD Biosciences, USA). Briefly, the biotin-SP-conjugated goat anti-mouse IgG F(ab’)2 fragment specific antibody was used as a primary antibody to capture CAR and the PerCP/Cy5.5 Streptavidin antibody (Biolegend, USA) was used as a secondary antibody. The CD19 expression levels are shown on Fig 2b. The sorted cells were cultured again to establish stable Jurkat/CAR T-cell lines.

### Transduction of lentivirus into hPBMCs

CD19 hPBMC CAR T-cells were generated by transduction of hPBMCs with lentivirus prepared as described elsewhere [7]. hPBMCs were drawn from healthy volunteer donors at the Research Institute of National Cancer Center following the Institutional Review Board-approved protocol.

### Synthesis of the ^89^Zr-DFO complex for cell labeling

^89^ZrCl_4_ was prepared from ^89^Zr-oxalate (Perkin Elmer). ^89^Zr-DFO complex was synthesized using 2.8 nmol of DFO and 62.9 ± 29.6 MBq of ^89^ZrCl_4_ and analyzed with a slight modification of the previously reported method [16]. We just changed the reagent of neutralization from KOH to NaOH and incubation time from 1 hr to overnight.

### Labeling of CAR T-cells with ^89^Zr-DFO

Cell labeling with ^89^Zr-DFO was performed with a slight modification of the previously described method [15]. Briefly, the Jurkat/CAR T-cells were counted, harvested and washed once with HBSS buffer (pH 7.5). Then, 185 kBq/100 μl HBSS buffer of ^89^Zr-DFO was added to each glass tube with 5 × 10^6^ cells in 500 ml HBSS buffer and incubated in a thermomixer at 37°C for 30 minutes (Eppendorf, Germany) with gentle shaking. After incubation, we added 1 ml of cold HBSS buffer, and the solution was centrifuged at with 1200 rpm for 5 minutes at 4°C in order to separate the supernatant. The supernatant was collected in a new tube and we repeated this washing process three times. The final cell labeling efficiency was calculated as follows: Labeling efficiency (%) = [cell activity (cpm)/ {cell activity (cpm) + supernatant activity (cpm)}6] × 100. CAR T-cells from hPBMC were labeled with the same procedure as the Jurkat /CAR T-cells, but ^89^Zr-DFO 74 kBq /100 μl HBSS buffer was added to each tube.

### Cell viability and proliferative activity of ^89^Zr-DFO-labeled cells

To investigate the cell viability and proliferation rates, ^89^Zr-DFO-labeled and unlabeled cells were seeded to 2 × 10^4^ ~ 5 × 10^5^ cells/well in 6 well plates containing 5 ml of culture medium. After seeding, the cell viability was assessed using trypan blue exclusion assay at 1 hour, 1 day, 3 days and 7 days. At the same time, the cell proliferation rate was compared with unlabeled cells. The seeded cells were maintained at 37°C, 5% CO_2_ incubator. Unlabeled cells served as controls.

### Function test for ^89^Zr-DFO-labeled cells

To determine the function of ^89^Zr-DFO-labeled Jurkat/CAR T-cells, we evaluated the target cell-specific cytokine IL-2 production ability with CD19 positive Raji cells (Burkitt lymphoma, ATCC, CCL-86) and CD19 negative K562 cells (chronic myelogenous leukemia, ATCC, CCL-243) as target cells. We then compared the results from ^89^Zr-DFO-labeled cells with the results obtained from unlabeled Jurkat/CAR T-cells. For IL-2 secretion assay, ^89^Zr-DFO-labeled cells were seeded 4 × 10^4^ cells/well in 6 well plates and were cultured for 12 or 36 hours at a 4:1 ratio [effector cells: target cells (Raji or K562 cells)]. Human IL-2 ELISA assay (RayBiotech, GA, USA) was performed according to the instructions of the manufacturer. To observe the function of ^89^Zr-DFO-labeled hPBMC CAR T-cells, we additionally performed IFN-γ release assay (RayBiotech, GA, USA) according to the instructions of the manufacture.

### Animal model establishment

To develop a mouse xenograft model, NSG mice (female, 5–6 weeks old, 20–22 g) from Jackson Laboratory (USA) were used. To compare tumor targeting of CAR T-cells, 5 × 10^6^ of Raji cells were injected into the left flank and 5 × 10^5^ of K562 cells were injected into the right flank of each mouse at the same time. Over the following 5–7 days, the tumors reached a volume of 50 to 100 mm^3^. Tumor volume was measured 3 times per week and calculated as length × width × height × π/6 (mm^3^).

### Animal PET-MR imaging and image analysis

PET-MRI fusion imaging was done using nanoScan PET/MRI (1T, Mediso, Hungary). ^89^Zr-DFO-labeled Jurkat/CAR T-cells (n, median: 4.1 × 10^6^, range: 3.1–5.4 × 10^6^; radioactivity, median: 907 kBq, range: 481–1221 kBq) were slowly injected intravenously through a tail vein using a 26 G syringe. Before PET image acquisition, the mice were kept under anesthesia (1.5% isoflurane in 100% O_2_ gas). Imaging was performed on days 0 (1 hour after the injection), 1, 2, 5, 6 and 7. The T1 weighted with gradient-echo 3D sequence (TR = 25 ms, TEeff = 3.4, FOV = 64 mm, matrix = 128×128) MR images were acquired, followed by static PET images for 10 minutes (days 0 and 1), 20 minutes (day 2), or 30 minutes (days 5, 6 and 7) in a 1:5 coincidence mode in a single field of view with the MRI range. Body temperature was controlled with heated air on the animal bedding (Multicell, Mediso, Hungry), and a pressure-sensitive pad was used for respiratory triggering. PET images were reconstructed using Tera-Tomo 3D, in full detector mode, with all the corrections on, high regularization and 8 iterations. A three-dimensional volume of interest (VOI) was applied to organs and tumors on the reconstructed PET and MR images using the InterView Fusion software package (Mediso, Hungary) and quantitative analysis procedures. Then, % injected dose (ID; radioactivity in each organ divided by injected radioactivity) were calculated. VOIs, with a fixed 2 mm diameter sphere, were also drawn for the tumors (Raji and K562), brains, hearts (left ventricle), lungs, livers, kidneys, spleens, and bones (femur), and they were analyzed using the following formula: standardized uptake value (SUV) = (radioactivity in the VOI with the unit of Bq/cc × body weight) divided by injected radioactivity.

### *Ex vivo* biodistribution study

Immediately after the PET-MR image acquisition on day 7, all mice were sacrificed. Their organs (brain, heart, lung, liver, spleen, kidney, stomach, intestine, bone, muscle, etc.) and tumors were excised, weighed, and counted by a gamma-counter for 5 minutes. The % ID values were obtained after normalization to the weight of each organ.

### Flow cytometry analysis

To validate the biodistribution of Jurkat/CAR T-cells after injection, we performed *ex vivo* immunostaining of organs. The mouse organs (liver, spleen) were harvested on day 3 after injection of unlabeled Jurkat/CAR T-cells (2 × 10^7^/200 μl) into the NSG mice with tumors via their tail veins (n=2). Tissues were ground using a gentleMACS™ dissociator (Miltenyi Biotec, Germany) according to the supplier’s protocol. A mouse cell depletion kit (Miltenyi Biotec, Germany) was used to separate the Jurkat/CAR T-cells from mouse cells. After counting the live cells, the human T-cell isolation kit (Miltenyi Biotec, Germany) was used according to the supplier’s protocol. Red blood cells were removed using a RBC lysing buffer (Sigma Aldrich, MO) for 1 minute, followed by washing and re-suspension in 1 x HBSS containing 1% FBS. The separated cells were used with PE-conjugated anti-CD3. The analysis staining process was the same as that used *in vitro*. Data were acquired from the stained cells using BD FACS CantoII flow cytometry (BD Biosciences). The results were evaluated with FlowJo software (Treestar Inc., Ashland, OR).

### Preparation of genomic DNA and Alu PCR analysis

To validate the biodistribution of Jurkat/CAR T-cells after injection, we also performed PCR analysis of *ex vivo* tissue (n = 3). At 3 days after injection of 2 × 10^7^ of Jurkat/CAR T-cells into the mice through the tail veins, the mice were sacrificed and brain, heart, lung, liver, spleen, and kidney samples were collected. For Alu PCR assay to detect injected human cells, genomic DNA was extracted from tissue samples using the QIAamp® (Qiagen, Germany) according to the protocol of the manufacturer. The primers used in this study were as follows: for human Alu, 5’-CACCTGTAATCCCAGCACTTT-3’ (forward primer) and 5’-CCCAGGCTGGAGTGCAGT-3’ (reverse primer). Real-time PCR was performed by SYBR® Green Realtime PCR Master Mix (TOYOBO, Japan) and the ABI 7500 Fast Real-Time PCR System (Applied Biosystems, CA, USA) according to the manufacturers’ instructions. The PCR experimental conditions were 95°C for 10 minutes, followed by 40 cycles at 95°C for 15 seconds and 60°C for 1 minute. This was followed by melting curve cycles at 95°C for 15 seconds, at 60°C for 1 minute, and finally at 95°C for 15 seconds. We used concentrations that were 1, 5, and 25 dilutions based on 10^6^ cells of Jurkat/CAR T-cells expressing CD19 as a standard. The results indicated the amount of Alu expression, based on the standard.

### Immunohistochemistry analysis

IHC analysis was performed as previously described [20]. The mice were sacrificed on day 3 (n = 3) or day 7 (n = 2) after Jurkat/CAR or hPBMC CAR T-cells were injected into tumor-bearing mice. For the day 3 group, after injection of Jurkat/CAR T-cells, the liver, spleen, and tumors were harvested, and fixed within paraffin blocks for IHC staining. The slides were stained with an anti-CD3 antibody (Abcam, UK) with the Dako REAL™ EnVision™ Detection System (Agilent Technologies, Inc., CA, USA) and were counter-stained with hematoxylin.

### Statistical analysis

Data are shown as mean ± SD unless otherwise stated. A value of p < 0.05 was considered statistically significant. The Kruskal-Wallis test was used to determine differences between time points after ^89^Zr-DFO labeling. And the Mann-Whitney test was used to determine differences between ^89^Zr-DFO-labeled and unlabeled cells. Statistical analyzes were performed by GraphPad Prism (GraphPad Software, CA, USA).

## Results

### Labeling efficiency, viability and proliferative ability of ^89^Zr-DFO-labeled Jurkat/CAR T-cells and CAR T-cells

The ^89^Zr-DFO labeling efficiency of Jurkat/CAR T-cells was 72.8% ± 11.0% at 185 kBq (Fig 3a). Cell viability after ^89^Zr-DFO labeling was 95.2% ± 1.2%, similar to levels obtained before labeling. The concentration of radioactivity for Jurkat/CAR T-cells was 103.6 kBq/10^6^ cells. The labeling of hPBMC CAR T-cells proceeded to 74 kBq. ^89^Zr-DFO labeling efficiency was similar to Jurkat/CAR T-cells (Fig 3a). The ^89^Zr-DFO-labeled hPBMC CAR T-cells showed a radioactivity concentration of 98.1 kBq/10^6^ cells.

**Fig 3.**
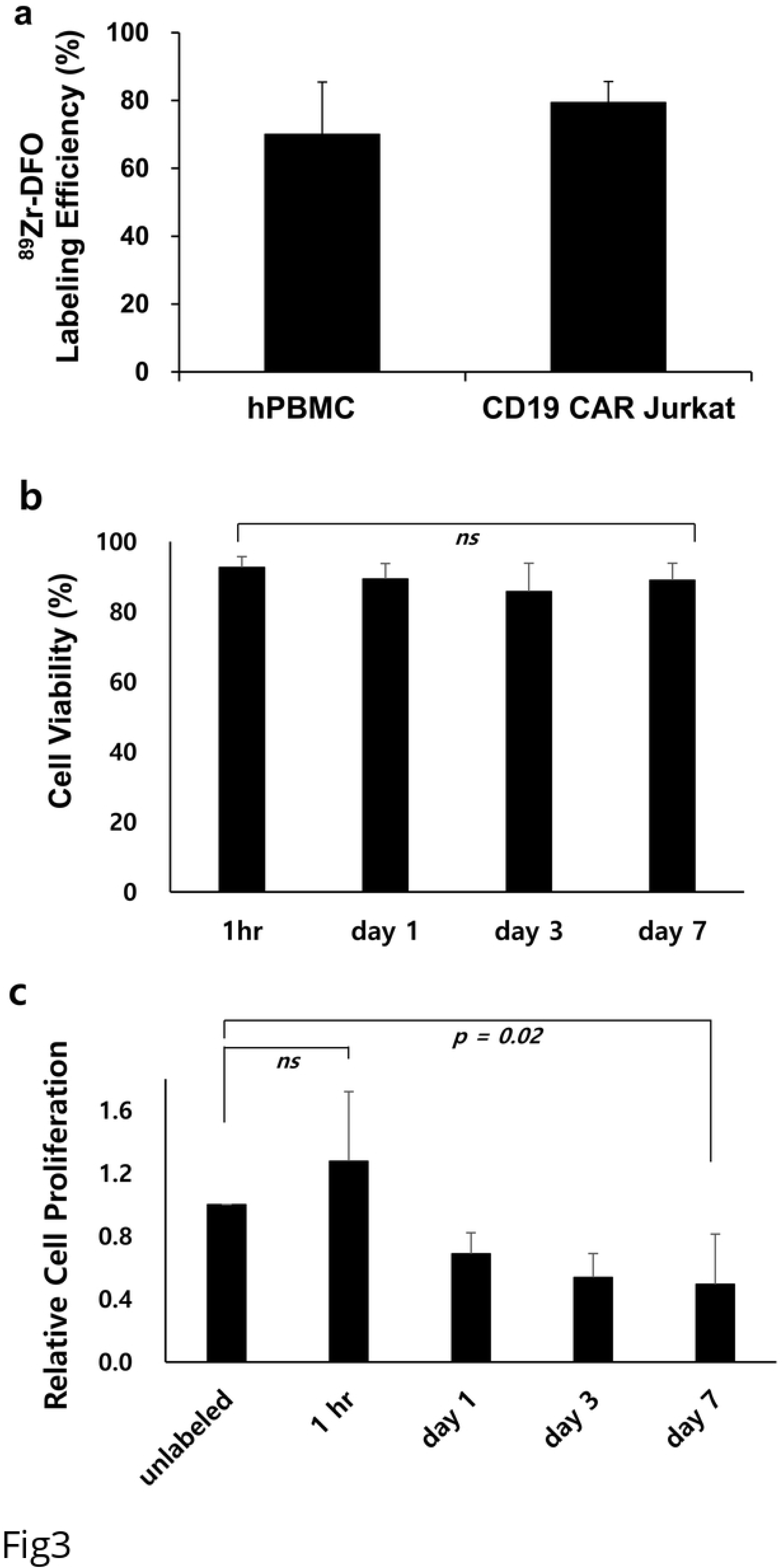
^89^Zr-DFO-labeled CAR T-cells labeling efficiency, cell viability and relative cell proliferation. **a** hPBMC CAR T-cells labeling efficiency after ^89^Zr-DFO labeling. **b** The cell viability after cell labeling was measured up to 7 days. **c** Relative cell proliferation of the labeled CAR T-cells was compared to that not labeled. Data are representative at least three independent experiments. All data are expressed as the mean and standard deviation.

We checked cell viability and cell proliferative ability at 1 hour, day 1, day 3 and day 7 after ^89^Zr-DFO cell labeling. There was no statistically significant difference in cell viability, although viability decreased over time after cell labeling (p = 0.24, 1 hour: 92.6% ± 3.1%, day 1: 89.4% ± 4.4%, day 3: 85.7% ± 8.1%, day 7: 89.0% ± 4.9%, respectively) (Fig 3b). However, the ^89^Zr-DFO-labeled cells showed decreased proliferative ability in a time-dependent manner (Fig 3c). At 1 hour after cell labeling, cell proliferation ability was not significantly changed, compared with unlabeled cells (*p* = 0.25). However, cell proliferation significantly decreased over time (1 hour: 1.28 ± 0.44, day 1: 0.69 ± 0.13, day 3: 0.54 ± 0.15, and day 7: 0.49 ± 0.32, respectively).

### Functional test for ^89^Zr-DFO-labeled Jurkat/CAR T-cells or CAR T-cells

CAR T-cell function was assessed by target cell-specific cytokine IL-2 production ability at 12 and 36 hours after cell labeling. Both ^89^Zr-DFO-labeled and unlabeled Jurkat/CAR T-cells induced IL-2 release in the positive target cells (Raji) at similar levels (74.1 vs 76.5 ng/ml at 12 hours, *p* = 0.99; 86.4 vs 87.1 ng/ml at 36 hours, *p* = 0.85) (Fig 4a). IL-2 was not released in negative target cells (K562) which did not react with Jurkat/CAR T-cells.

**Fig 4.**
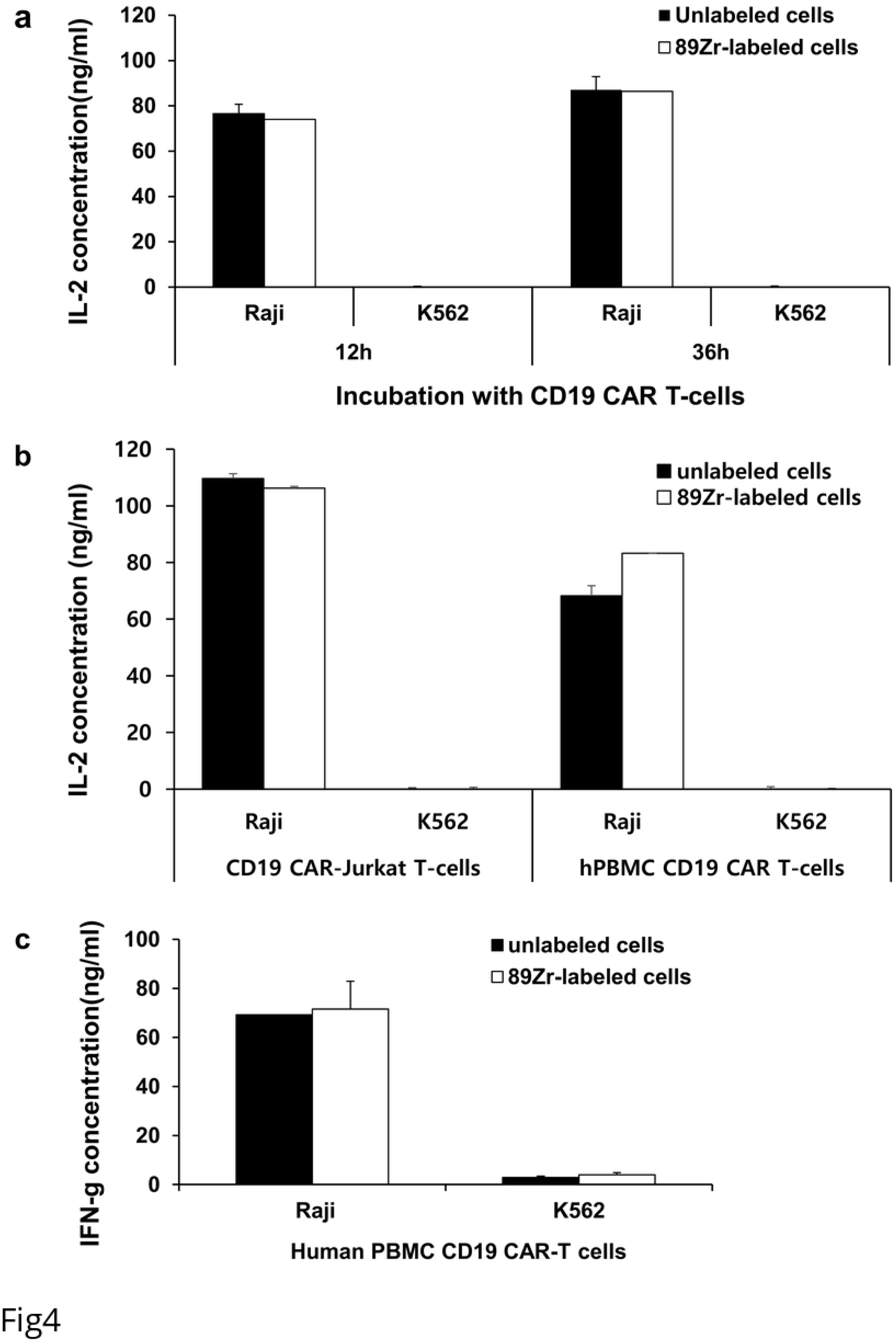
Cell function test after ^89^Zr-DFO-labeled Jurkat/CAR T-cells and hPBMC CAR T-cells. **a** Function test by IL-2 production of ^89^Zr-DFO-labeled Jurkat/CAR T-cells after incubation with Raji (CD19 positive) or K562 (CD19 negative) cells. Incubation time was for 12 hours or 36 hours and unlabeled cells were used in the experiment as a control group. **b** Function test by IL-2 production of ^89^Zr-DFO-labeled Jurkat/CAR T-cells or hPBMC CAR T-cells after incubation with Raji or K562 cells. **c** IFN-γ test of ^89^Zr-DFO-labeled hPBMC CAR T-cells with Raji or K562 cells. Data are representative at least two independent experiments. All data expressed as means and standard deviations.

The IL-2 secretion of hPBMC ^89^Zr-DFO-labeled CAR T-cells was also similar with that observed in unlabeled cells (Fig 4b). When hPBMC CAR T-cells were added to Raji cells expressing the targeted CD19 cells, the secretion of IFN-γ involved in cell cytotoxicity was maintained after ^89^Zr-DFO labeling (69.3 vs 71.6 ng/ml, *p* = 0.67) (Fig 4c).

### Trafficking of ^89^Zr-DFO-labeled Jurkat/CAR T-cells with PET/MR imaging

After injection of ^89^Zr-DFO-labeled Jurkat/CAR T-cells through the tail veins of the mice, we used PET/MR images for noninvasive real-time tracking of injected cells *in vivo*. The Jurkat/CAR T-cells were initially located in the lungs, then redistributed to the liver and spleen (Fig 5a). In more detail, the injected ^89^Zr-DFO-labeled Jurkat/CAR T-cells were found mainly in the lung (24.4% ± 3.4%ID) and liver (22.9% ± 5.6%ID) during the first 1 hour. Over time, the CAR T-cells gradually migrated from the lungs and accumulated mainly in the liver, some in spleen (Fig 5b). However, radioactivity accumulation was not evident in either CD19-positive Raji or CD19-negative K562 tumors. The detailed quantitative values of the PET images analyzed by SUV are shown in Table 1. Immediately after PET/MR imaging on day 7, the mice were sacrificed, then their organs and tumors were isolated to measure the radioactivity of injected cells *ex vivo*. Biodistribution data measured *ex vivo* on day 7 showed similar patterns of distribution assessed by PET images as shown in Fig 5c and Table 2.

**Table 1.** Quantitative analyses of ^89^Zr-DFO-labeled Jurkat/CAR T-cells on PET images with SUV. (mean ± SD, n = 4)

| Tissues | Day 0 (1 hour) | Day 1 | Day 2 | Day 5 | Day 6 | Day 7 |
| --- | --- | --- | --- | --- | --- | --- |
| Raji | $0.29 \pm 0.04$ | $0.26 \pm 0.04$ | $0.25 \pm 0.05$ | $0.21 \pm 0.01$ | $0.19 \pm 0.00$ | $0.20 \pm 0.03$ |
| K562 | $0.51 \pm 0.18$ | $0.43 \pm 0.09$ | $0.53 \pm 0.22$ | $0.42 \pm 0.11$ | $0.37 \pm 0.04$ | $0.35 \pm 0.10$ |
| Brain | $0.05 \pm 0.01$ | $0.09 \pm 0.01$ | $0.10 \pm 0.01$ | $0.12 \pm 0.02$ | $0.13 \pm 0.02$ | $0.15 \pm 0.02$ |
| Heart | $1.19 \pm 1.16$ | $0.39 \pm 0.03$ | $0.36 \pm 0.07$ | $0.23 \pm 0.04$ | $0.26 \pm 0.07$ | $0.24 \pm 0.07$ |
| Lung | $11.72 \pm 1.84$ | $4.31 \pm 0.67$ | $3.50 \pm 0.40$ | $2.63 \pm 0.49$ | $2.67 \pm 0.60$ | $2.22 \pm 0.46$ |
| Liver | $7.02 \pm 0.69$ | $9.40 \pm 0.60$ | $10.24 \pm 0.86$ | $10.16 \pm 0.69$ | $10.28 \pm 0.83$ | $9.54 \pm 0.51$ |
| Spleen | $5.29 \pm 0.49$ | $7.71 \pm 0.99$ | $9.02 \pm 2.78$ | $10.59 \pm 0.89$ | $9.81 \pm 1.22$ | $10.22 \pm 0.59$ |
| Kidney | $0.36 \pm 0.04$ | $0.35 \pm 0.04$ | $0.39 \pm 0.05$ | $0.49 \pm 0.08$ | $0.42 \pm 0.04$ | $0.51 \pm 0.09$ |
| Bone | $0.23 \pm 0.04$ | $0.79 \pm 0.23$ | $0.81 \pm 0.36$ | $0.98 \pm 0.19$ | $0.88 \pm 0.17$ | $0.81 \pm 0.16$ |

**Table 2.** Quantitative analyses of *ex vivo* biodistribution of ^89^Zr-DFO-labeled Jurkat/CAR T-cells study on day 7. (%ID/g; mean ± SD, n = 4)

| Tissues | %ID/g |
| --- | --- |
| Raji | 0.09 ± 0.01 |
| K562 | 0.08 ± 0.01 |
| Blood | 0.04 ± 0.01 |
| Brain | 0.01 ± 0.01 |
| Heart | 0.12 ± 0.05 |
| Lung | 24.01 ± 8.05 |
| Liver | 27.01 ± 2.96 |
| Spleen | 118.65 ± 87.60 |
| Kidney | 0.30 ± 0.10 |
| Stomach | 0.06 ± 0.01 |
| S-intestine | 0.04 ± 0.01 |
| L-intestine | 0.03 ± 0.01 |
| Ovary | 0.26 ± 0.25 |
| T-spine | 1.13 ± 0.17 |
| Femur | 2.31 ± 1.06 |
| Muscle | 0.04 ± 0.01 |

**Fig 5.**
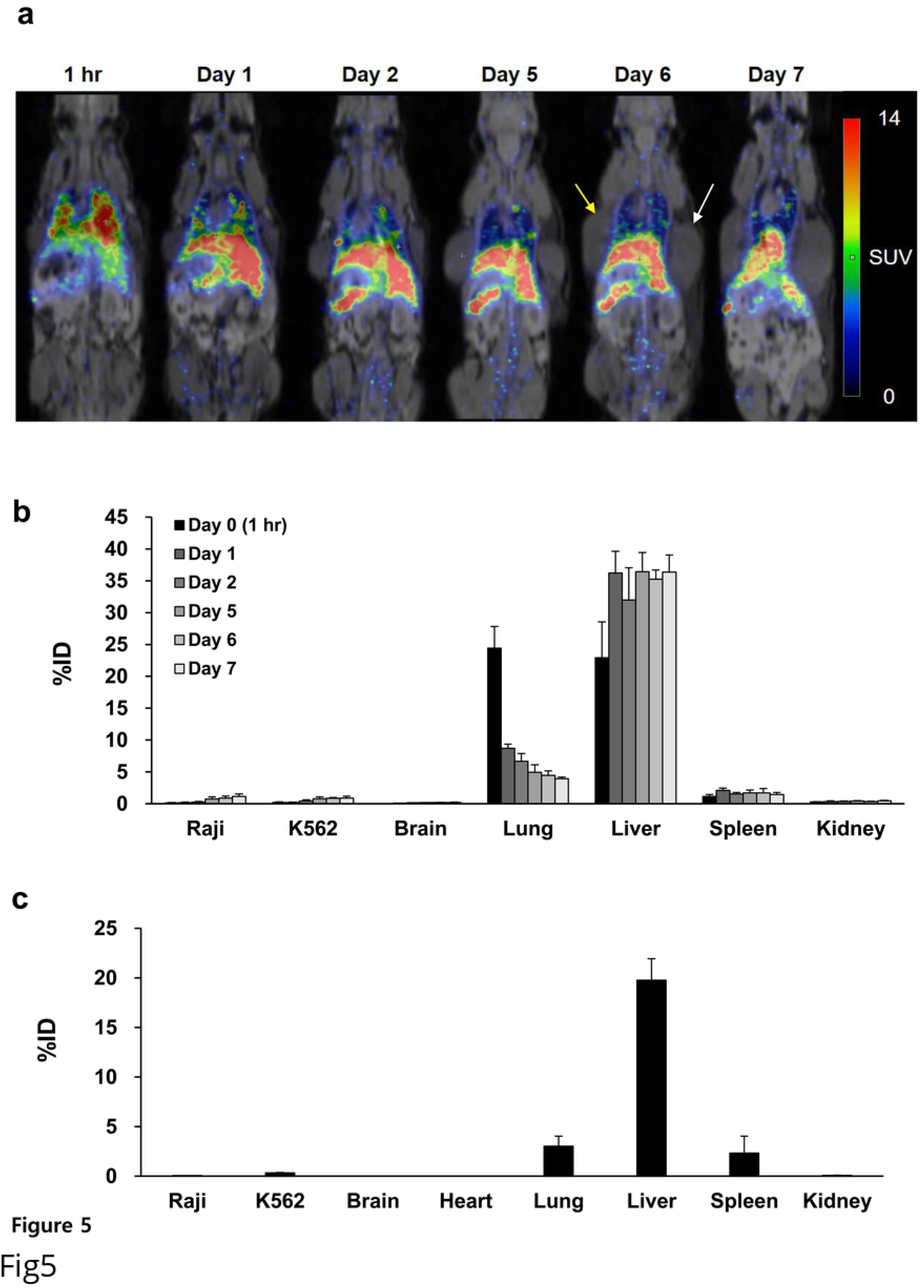
Serial ^89^Zr-DFO-labeled Jurkat/CAR T-cells animal PET/MR images and analysis of biodistribution. **a** PET images of the whole-body distribution of intravenously injected ^89^Zr-DFO-labeled CAR T-cells in NSG mouse xenograft following until day 7. Yellow arrow and white arrow represent Raji tumor and K562 tumor, respectively. **b** The distribution of *in vivo* organ measured by ^89^Zr-DFO-labeled Jurkat/CAR T-cells PET imaging over the following 7 days. **c** The distribution of *ex vivo* organ measured by model sacrifice after acquisition of the last PET imaging. (n = 4)

PET/MR imaging with ^89^Zr-DFO-labeled CAR T-cells from hPBMC also showed similar biodistribution of cells after tail vein injection. We did not observe increased radioactivity in the tumors, which would have suggested CAR T-cell homing (Fig 6). The detailed quantitative values analyzed by SUV are shown in Table 3.

**Table 3.** Quantitative analyses of ^89^Zr-DFO-labeled hPBMC CAR T-cells on PET images with SUV. (mean ± SD, n = 2)

| Tissues | Day 0 (1 hour) | Day 0 (4 hours) | Day 1 | Day 3 | Day 6 |
| --- | --- | --- | --- | --- | --- |
| Raji | 0.57 ± 0.12 | 0.74 ± 0.45 | 0.55 ± 0.16 | 0.34 ± 0.12 | 0.47 ± 0.23 |
| K562 | 0.14 ± 0.02 | 0.20 ± 0.11 | 0.20 ± 0.03 | 0.18 ± 0.00 | 0.29 ± 0.07 |
| Brain | 0.10 ± 0.01 | 0.12 ± 0.02 | 0.12 ± 0.04 | 0.15 ± 0.04 | 0.51 ± 0.43 |
| Heart | 0.74 ± 0.37 | 0.56 ± 0.20 | 0.27 ± 0.14 | 0.32 ± 0.04 | 0.32 ± 0.02 |
| Lung | 5.79 ± 0.25 | 3.71 ± 0.03 | 2.18 ± 0.33 | 2.15 ± 0.20 | 1.99 ± 0.63 |
| Liver | 9.07 ± 0.04 | 10.51 ± 1.23 | 10.20 ± 0.31 | 10.38 ± 0.16 | 8.81 ± 3.19 |
| Spleen | 4.34 ± 0.32 | 7.29 ± 0.88 | 6.12 ± 0.37 | 3.78 ± 0.27 | 4.30 ± 1.09 |
| Kidney | 0.81 ± 0.50 | 0.62 ± 0.15 | 0.43 ± 0.09 | 0.54 ± 0.00 | 1.22 ± 0.61 |
| Bone | 0.30 ± 0.15 | 0.49 ± 0.14 | 0.64 ± 0.35 | 0.48 ± 0.40 | 1.19 ± 0.71 |

**Fig 6.**
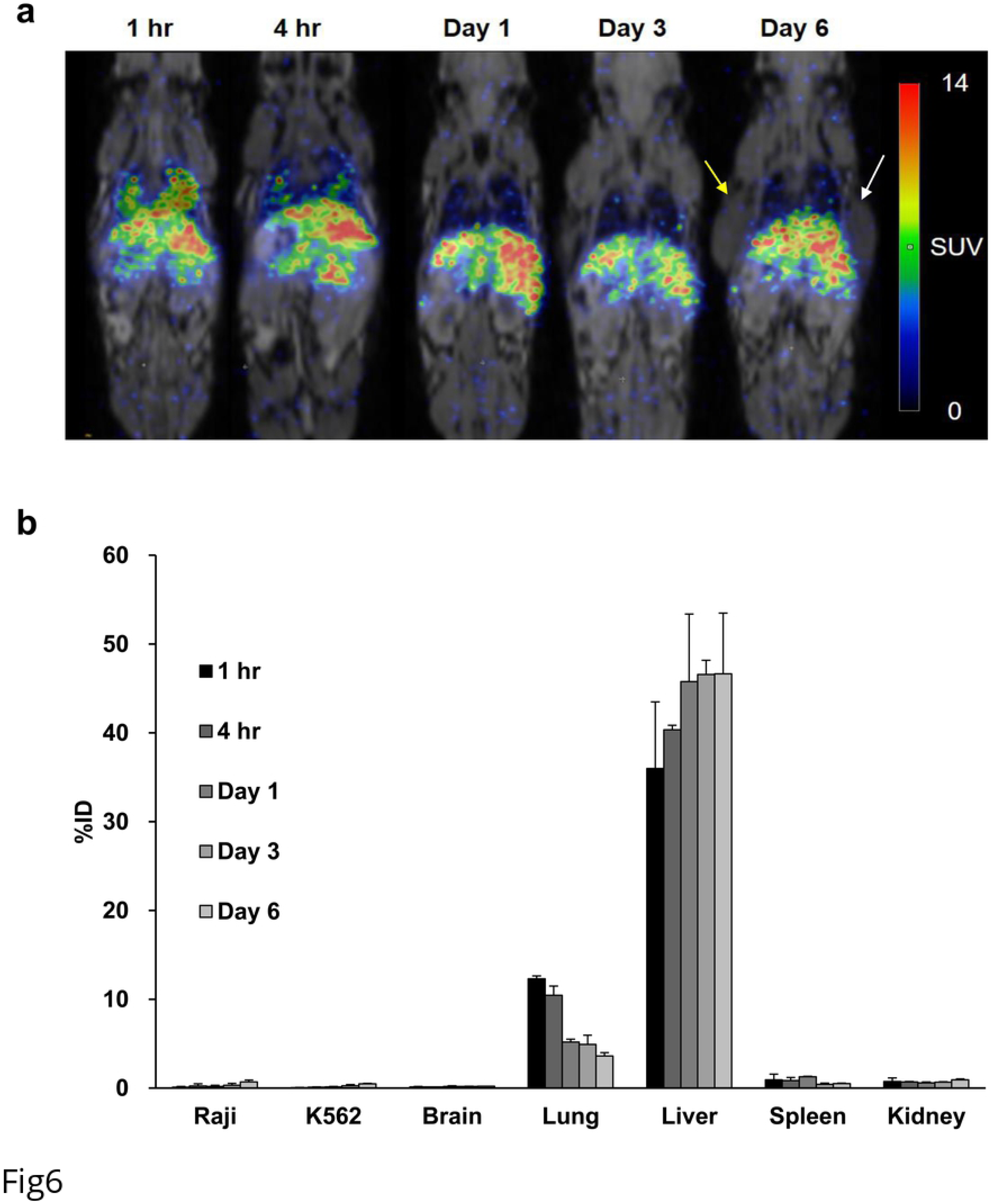
Serial ^89^Zr-DFO-labeled hPBMC CAR T-cells animal PET/MR images and analysis of biodistribution. **a** PET images of whole-body distribution for intravenously injected ^89^Zr-DFO-labeled hPBMC CAR T-cells in NSG mouse xenograft following until day 6. Yellow arrow and white arrow represent Raji tumor and K562 tumor, respectively. **b** The distribution of *in vivo* organ measured by ^89^Zr-DFO PET imaging during 6 days. (n = 2)

### Flow cytometry, Alu PCR and immunohistochemistry for *ex vivo* organ study

We evaluated biodistribution after injection of unlabeled Jurkat/CAR T-cells using non-imaging methods, for the comparison with using imaging method. For flow cytometry, the liver and spleen tissue samples from mice were separated into single cells. After the initial separation an average number of 3.05 × 10^7^ and 1.67 × 10^6^ cells per mouse were harvested. After depletion of mouse liver cells, an average of 4.74 × 10^4^ cells was obtained after using a human T-cell isolation kit. In contrast to the negative control stained with isoform antibody, CD3 expressing CAR T-cells were distributed in 95.7% of the spleen and 60.3% of the liver (Fig 7a).

**Fig 7.**
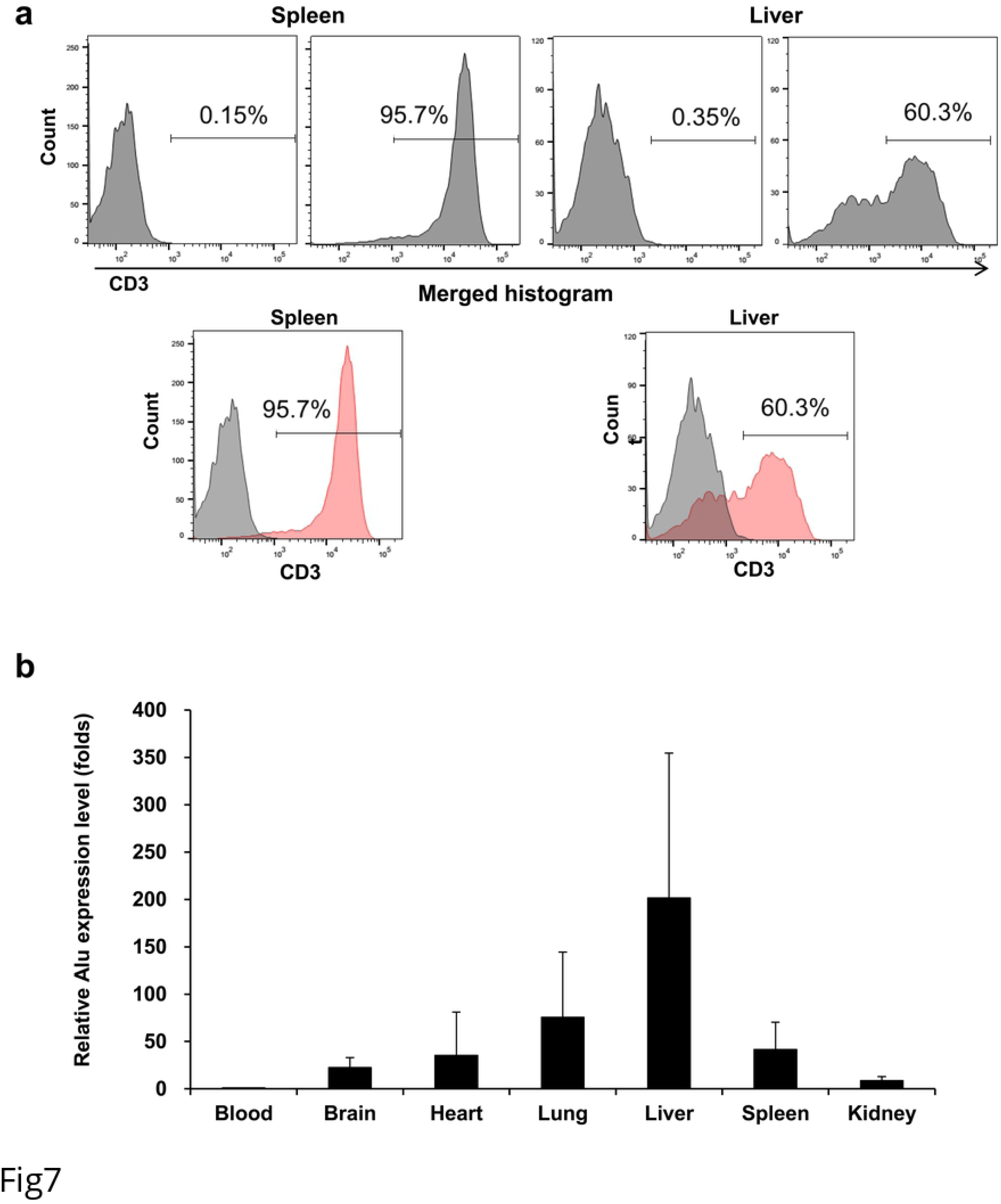
FACS staining and Alu PCR in mouse organs. **a** Post sacrificing the mice on day 3 after Jurkat/CAR T-cells injection, graphs of FACS staining for liver and spleen tissues of mice were plotted against the control group. **b** Alu PCR analysis data of mouse blood, brain, heart, lung, liver, spleen, kidney and gut tissues obtained from sacrifice 3 days after Jurkat/CAR T-cells injection.

Organ distributions of Jurkat/CAR T-cells were confirmed through Alu PCR. The blood, heart, lung, liver, spleen and kidney were sampled, and CD19 CAR T-cells were distributed in all 6 organs. The relative Alu expression was determined by measuring blood, and the fold was as follows: brain (15.1 ± 5.8), heart (6.2 ± 7.2), lung (38.5 ± 34.8), liver (212.2 ± 225.4), spleen (70.3 ± 9.4), kidney (9.4 ± 6.4) and gut (17.5 ± 8.9) (Fig 7b).

Immunohistochemical staining with CD3 of liver and spleen tissues confirmed the presence of Jurkat/CAR T-cells in the liver and spleen, in contrast to control tissue from mice not injected with Jurkat/CAR T-cells (Fig 8a). Immunohistochemical staining of Raji and K562 tumors of the day 3 group with CD3 showed barely visible stained cells in the periphery of the tumors. In the day 7 group, hPBMC CAR T-cells showed proliferation throughout Raji tumors; however, Jurkat/CAR T-cells were not visualized (Fig 8b). No K562 tumors were stained with CD3 antibody.

**Fig 8.**
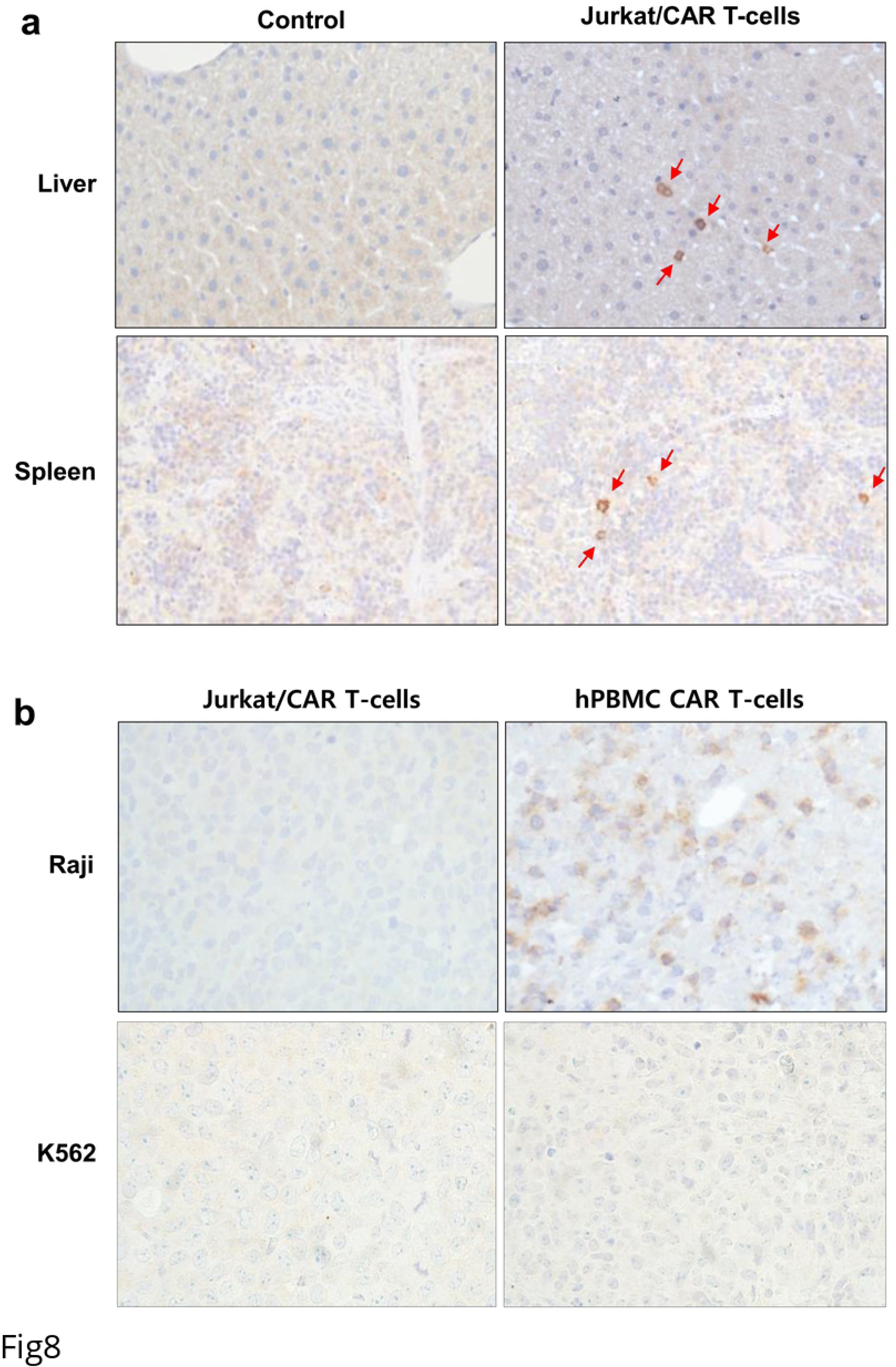
IHC staining in mouse organs. **a** IHC staining with CD3 antibody demonstrates increased staining in liver and spleen tissue after Jurkat/CAR T-cells were injected into a mouse, compared to control mouse spleen. Red arrows show CD3 targeting T-cells in IHC staining. **b** IHC staining with CD3 antibody in Raji and K562 tumor tissues on day 7 after Jurkat/CAR or hPBMC CAR T-cells injection.

## Discussion

In this study, *in vivo* CAR T-cell trafficking was feasible for 7 days after intravenous injection by ^89^Zr-DFO labeling and PET/MR images. CAR T-cells initially reached the lungs and gradually migrated to the liver by day 1, where they remained for the rest of the experimental period. Migration to the spleen was also evident and showing high SUV on PET/MR images, although %ID was relatively low and size of the spleen is small in immunocompromised NSG mice. The organ distribution of ^89^Zr-DFO-labeled cells quantitatively assessed by PET/MR images was confirmed with an *ex vivo* biodistribution study where we analyzed the radioactivity of each organ harvested from the mice on day 7. This pattern of distribution was observed in both ^89^Zr-DFO labeling of Jurkat/CAR T-cells and CAR T-cells from human peripheral blood. The distribution of cells after injection of unlabeled Jurkat/CAR T-cells was also confirmed by flow cytometry, Alu PCR, and IHC with isolated tissues from sacrificed mice on day 3. We could reliably and noninvasively track the distribution after cell administration using ^89^Zr-DFO labeling of CAR T-cells and PET/MR imaging, as suggested by FDA guidance. In quantitative analysis of organ distribution of cell populations, previous studies with radioisotope labeling had a lack of specific organ distribution data because of the difficulties in gathering anatomical information with PET alone. The current study was performed using a hybrid imaging system. Here PET and MR allowed organ-specific detection of the cell signal, along with structural information. The quantitative nature of PET allows for longitudinal studies that provide information on relative levels of CAR T-cells at both the site of disease and potential off-target sites of accumulation. Monitoring the locations and potential secondary sites involved in CAR T-cell trafficking enables us to characterize the activity of administered cells and safety profile.

^89^Zr-DFO was used for labeling CAR T-cells in this study, instead of ^89^Zr-oxine complex, which was used by Weist et al. [19], because it has more stable covalent binding between DFO and cell surface protein [16]. With ^89^Zr-DFO labeling of human immune cells, radioactivity concentrations of radioisotope-labeled cells of up to 0.5 MBq/10^6^ cells were executed without an unfavorable effect on cellular viability and cell efflux studies showed high radiolabel stability, with virtually no loss of tracer for up to 7 days [16]. Whereas a significant amount of ^89^Zr-oxine can efflux from various kind of cells, and there is potential uptake of small amount of free ^89^Zr released from cells into the bones or kidneys, as shown in the report by Weist et al. [19]. In our PET images of ^89^Zr-DFO-labeled cells, less radioactivity accumulated in the bones and kidneys until 7 days after injection, compared with the results reported with ^89^Zr-oxine labeled cells. Furthermore, decay corrected injected activity was maintained on serial PET images (98.9 ± 8.3%) without significant loss of activity from the body. These findings suggest stable cell labeling by ^89^Zr-DFO *in vivo*, as demonstrated in other studies.

The cell viability and functionality such as cytokine production were not affected by labeling CAR T-cells with ^89^Zr-DFO, at radioactivity cell concentration of 98 kBq/10^6^ cells. However, proliferation ability was slightly decreased over the next several days compared with unlabeled cells. It is important to consider the effect of radiation dose on cells of hematopoietic origin, which are relatively radiation sensitive. Especially, ^89^Zr has high-energy gamma emissions of 908.97 keV, which may limit the radioactive dose; further, the radiolabeling procedure is potentially cytotoxic. Therefore, we reduced the dose of ^89^Zr-DFO when labeling CAR T-cells from hPBMC to 74 kBq/10^6^ cells, compared with 185 kBq/10^6^ cells in the labeling of Jurkat/CAR after careful dose optimization because of hPBMC sensitivity.

The distribution pattern of CAR T-cells in this study was similar with previous studies. Following intravenous administration, human T-cells migrated in a manner similar to that reported in humans, but penetrated poorly into established tumors. Following intravenous administration, human T-cells initially reached the lungs where they remained for more than 4 hours. After that, the T-cells redistributed to the liver, spleen, and lymph nodes [20]. This pattern was seen using all T-cell populations tested, regardless of tumor status or transgene cargo, and closely mimicked patterns of migration seen in human (via infusion) [21–23] or murine T-cell [24,25] recipients. According to Charoenphun et al., ^89^Zr-oxine labeled myeloma cells were injected intravenously and were found within the lungs at 30 minutes after injection, but migrated to the liver and spleen on day 1. This distribution continued until day 7 [26]. Similar distributions of ^89^Zr -oxine labeled dendritic cells were observed by Sato et al. [18]. Weist et al. also found that the highest CAR T-cell activity in the spleen, followed by the liver [19]. In these studies, lung activity was significantly lower on day 7 than activity in the liver or spleen. In contrast, Bansal et al. found that mesenchymal stem cells exhibited persistently high activity in lung images, up to day 7, and their biodistribution study also showed the highest activity in the lung (approximately 50%ID), followed by the liver (approximately 25%ID) [16]. It has been suggested that the slow migration of transferred T-cells through the lungs may be due to low pulmonary circulatory pressure, coupled with the narrowing of capillaries during expiration [23]. Importantly, activated T-cells cross the pulmonary circulation with reduced intravascular velocity, compared to their inactivated or naive counterparts [27,28]. This delay may reflect an enhanced interaction between high-affinity state LFA-1 (T-cells) and ICAM-1 (endothelium), in part [27]. Delayed clearance of activated T-cells during their first pass through the lungs may be highly related with pulmonary toxicity that can occur following infusion of CAR T-cells. In one published incident, fatal adult respiratory distress syndrome occurred rapidly following infusion of >10^10^ T-cells, targeted against ErbB2 using a trastuzumab scFv coupled to a fused CD28/4-1BB/CD3ζ endoplasmic domain [29].

In this study, T-cell migration to the target tumor was not observed on PET images, unlike reports by Sato et al. and Weist et al. It was disappointing to observe that only a minority of intravenously administered CAR T-cells migrated to tumor deposits, even though we used CD19 CAR T-cells with proven efficacy in animals and humans. There are several potential causes of this phenomenon. We used Jurkat/CAR T-cells in this study are leukemia cells, and not true T-cells. Although Jurkat cells were used for convenience in this experiment, their actual biologic behavior may be different from normal T-cells. Second, the solid subcutaneous tumor xenograft model used in this study was a different tumor environment from that of human acute lymphocytic leukemia of the blood, for which CD19-CAR T-cells have shown a dramatic therapeutic effect. Also, the differences in the antigen-presenting status of the tumor, tumor microenvironment etc. between the xenograft model generated from Raji cells and those of previous studies by Sato et al. (B16 murine melanoma cells [18]) and Weist et al. (prostate cancer PC3-overexpressed cells [19]) may have affected the accumulation of CAR-T-cells within tumor tissue [30]. Third, detection of cell trafficking by imaging carries important limitations. Cell trafficking may not be detectable via PET imaging when small amounts of T-cells, below detectable limits, are injected. Furthermore, after homing to the tumor, activated CAR T-cells can proliferate and dilute the labeling signal. In our IHC & PET imaging data, hPBMC CAR T-cells showed proliferation within Raji tumor tissues after CD19 targeting (Fig 8b); however, there are rare PET imaging signals in tumors.

There are several limitations to this study. First, we did not show a therapeutic effect for CAR T-cells, because our experiments were conducted mainly with Jurkat/CAR T-cells instead of hPBMC CAR T-cells. Limited imaging experiments were possible with hPBMC CAR T-cells and only two mice. Second, the biodistribution study was only performed using direct cell labeling strategies. The direct cell labeling method cannot visualize cell proliferation after homing to the target tumor [31].

In this study, ^89^Zr-DFO labeling CAR T-cells direct imaging had great advantages to be able to observe the initial behavior of the injected cells and the distribution in the whole body in real-time with very low background activities which is in principle not present in other host tissues or cells. And this direct labeling did not involve genetic manipulation of the therapeutic cells therefore it was a simple way to trace initial injected CAR T-cells.

Further studies that use both direct and indirect labeling strategies, where reporter genes are inserted in the vector, should be carried out with hPBMC CAR T-cells.

## Conclusion

With ^89^Zr-DFO labeling of CAR T-cell, real-time *in vivo* cell trafficking was feasible through PET imaging after administration of cells to the body. Thus, ^89^Zr-DFO-labeled CAR T-cell PET imaging can be used to investigate cell kinetics, *in vivo* cell biodistribution, and the safety profile of CAR T-cell therapies that are developed.

## Supporting Information

S1 File. Original data of the present study

## Acknowledgements

Not applicable.

